## Supplementary figures and table for "Role of the *osaA* gene in *Aspergillus fumigatus* development, secondary metabolism and virulence"

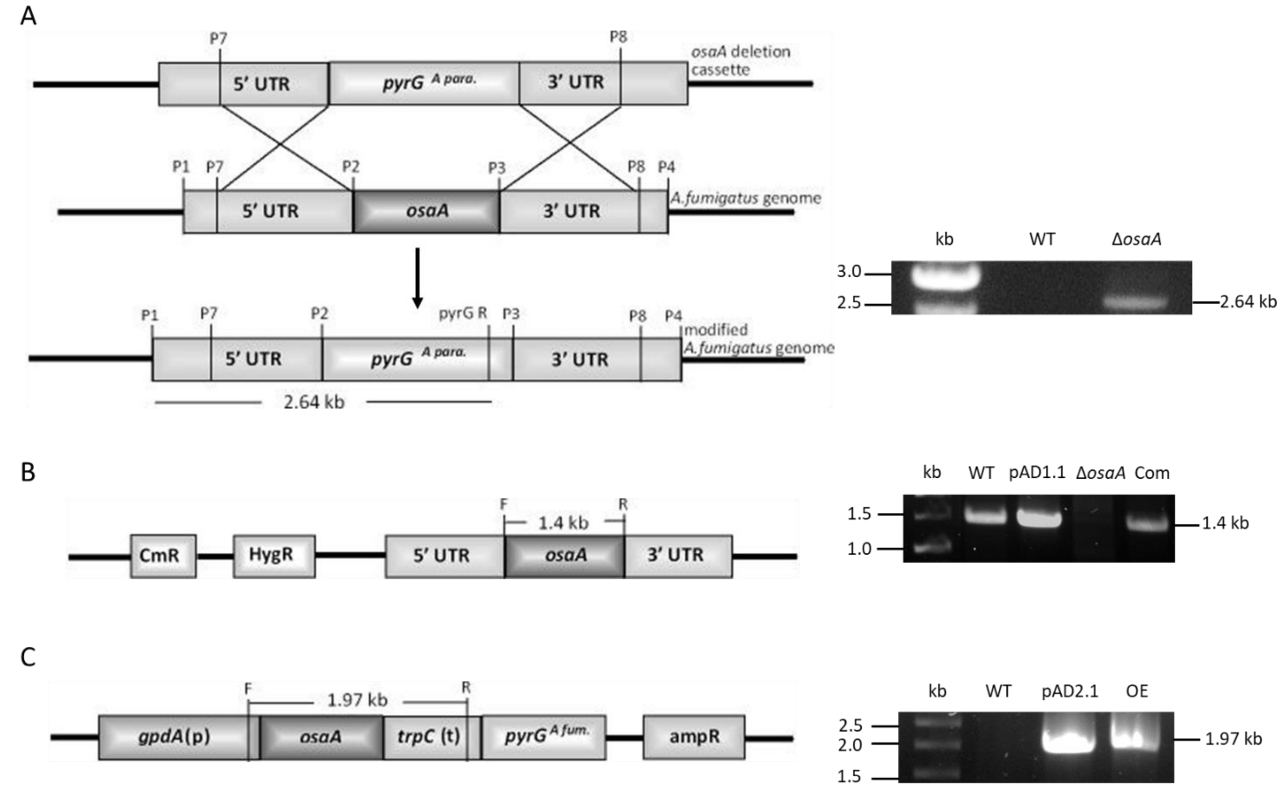


**Figure S1. Generation of the *osaA* deletion, complementation and overexpression strains.** (A) Generation of the *osaA* deletion strain (Δ*osaA*). Diagram showing the replacement of *osaA* gene with the *A. parasiticus pyrG* marker gene by homologous recombination. On the right, diagnostic PCR using primers P1 (osaA_F1) and pyrGR (A para pyrG_R) resulting in the expected 2.64 kb PCR product. Wild-type strain was used as negative control. (B) Generation of the *osaA* complementation strain (Com). Schematic representation of *osaA* complementation plasmid (pAD1.1). The hygromycin resistance gene, HygR, was used as a transformation marker. On the right, confirmation of the complementation transformant by diagnostic PCR using primers *osaA*F (osaA_OE1) and *osaA*R (osaA_OE2). The expected PCR product of 1.4 kb was observed. Wild type and pAD1.1 were used as positive controls, *osaA* deletion strain was used as negative control. (C) Generation of the *osaA* overexpression strain (OE). Representation of the *osaA* complementation plasmid (pAD2.1). On the right, diagnostic PCR confirmation of *osaA* OE strain, showing the expected 1.97 kb product amplified with primers *gpdA*F (gpdA_F) to *trpC* (trpC_R). pAD2.1 was used as positive control, and wild type was used as negative control.

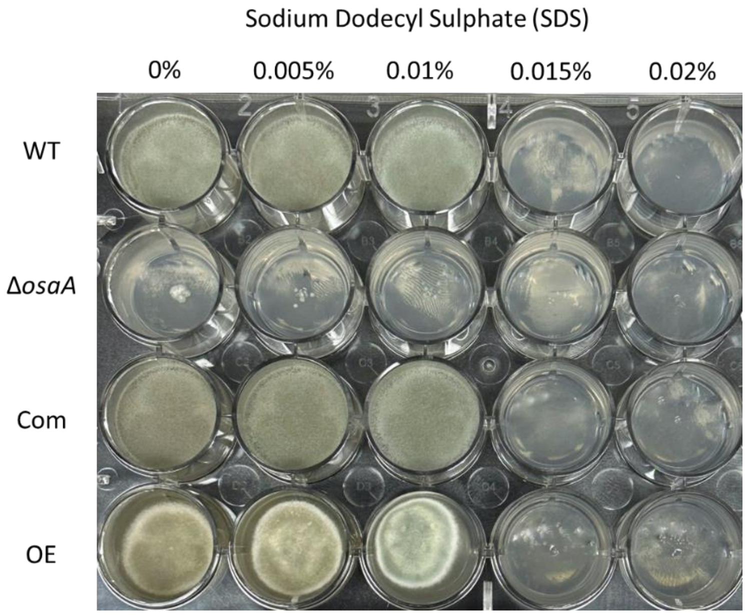


**Figure S2. Deletion *osaA* is more susceptible than wild type to the presence of sodium dodecyl sulphate (SDS).** *A. fumigatus* strains were point-inoculated on GMM and GMM supplemented with a range of sodium SDS concentrations. The plates were incubated at 37°C for 5 days. The experiment was performed in triplicates.

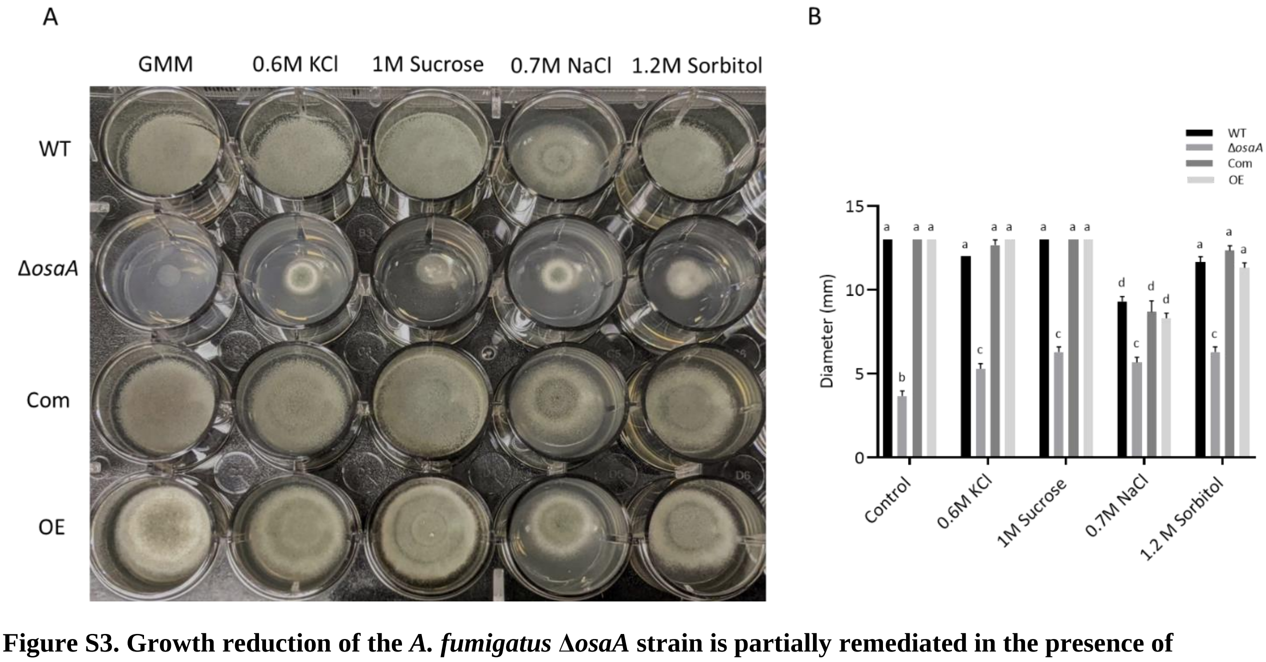


**Figure S3. Growth reduction of the *A. fumigatus* Δ*osaA* strain is partially remediated in the presence of osmotic stabilizers.** (A) *A. fumigatus* wild type (WT), deletion *osaA* (Δ*osaA*), complementation (Com) and overexpression (OE) strains were point-inoculated on GMM and GMM supplemented with 0.6 M KCl, 1 M sucrose, 0.7 M NaCl or 1.2 M sorbitol. Plates were incubated at 37°C for 72 h. The experiment was carried out in triplicate. (B) Colony diameter measurement of *A. fumigatus* cultures. Error bars indicate the standard error. Different letters on the columns indicate values that are statistically different ($P < 0.05$). All the experiments were done in triplicates.

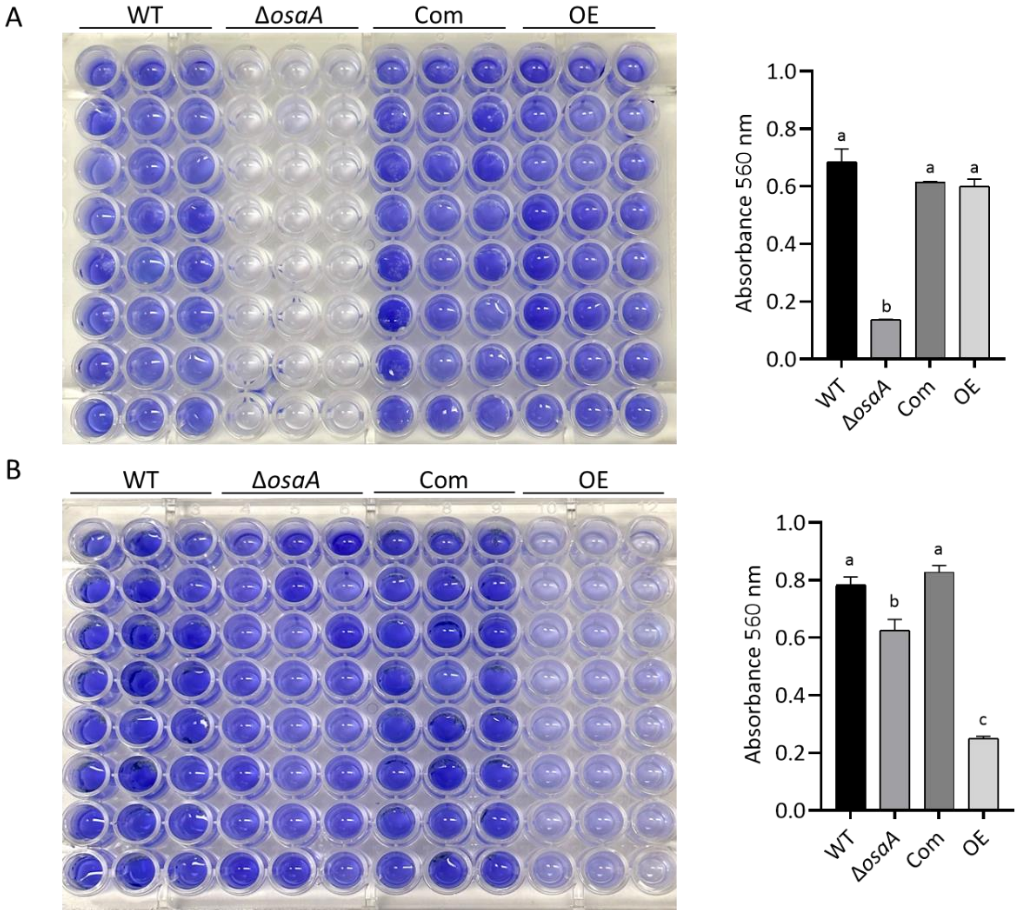


**Figure S4. *osaA* is necessary for normal adhesion to solid surfaces in *A. fumigatus*.** Wild type (WT), Δ*osaA*, complementation (Com) and overexpression (OE) strains were grown in 96-well plates containing 130 µL liquid GMM at 37°C. Twenty-four replicates were considered per strain. Adherence was measured as described in the Materials and Methods section. The bar graphs represent absorbance readings at 560 nm after (A) 24 h and (B) 48 h. Error bars indicate the standard error. Different letters on the columns indicate values that are statistically different ($P < 0.05$).

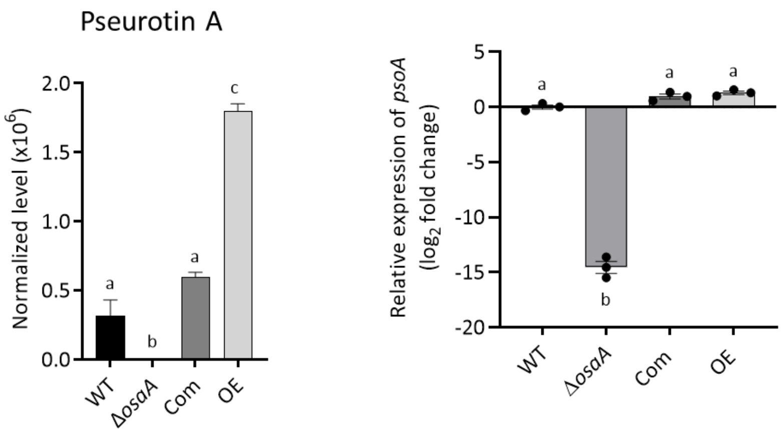


**Figure S5. *osaA* affects production of pseurotin A in *A. fumigatus*.** Wild type (WT), Δ*osaA*, complementation (Com) and overexpression (OE) strains were top-agar inoculated on GMM and incubated at 37°C for 72 h. Extracts were analyzed by liquid chromatography coupled to high resolution mass spectrometry (LC-HRMS). On the right, gene expression of *psoA*, a gene involved in pseurotin A production, was analyzed using the $2^{-\Delta\Delta CT}$ method [86]. WT, Δ*osaA*, Com and OE strains were grown in GMM liquid stationary cultures at 37°C for 48 h. Error bars indicate the standard error. Different letters on the columns indicate values that are statistically different ($P < 0.05$). All the experiments were done in triplicates.

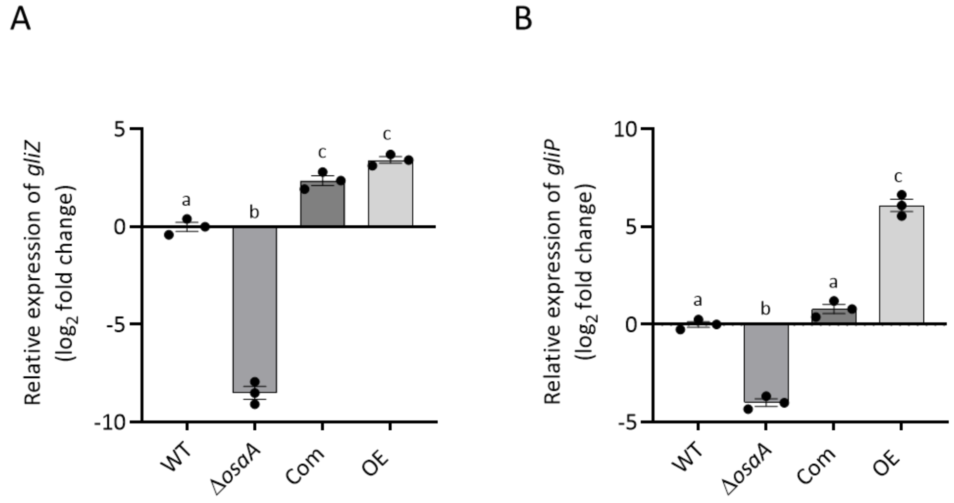


**Figure S6. Gene expression of *gliZ* (A) and *gliP* (B) is *osaA*-dependent**. Wild type (WT), Δ*osaA*, complementation (Com) and overexpression (OE) strains were grown in liquid shaken YES medium for 72 h at 250 rpm. Expression of genes were analyzed by qRT-PCR and relative expression was calculated following the $2^{-\Delta\Delta CT}$ method [86]. Tubulin gene expression was used as internal reference. Error bars represents standard error. Different letters on the bar represents significantly different values ($P < 0.05$).

**Table S1. Primers used in this study.**

| Name | 5'-3' sequence |
|---|---|
| 1. osaA_F1 | CCAGGTTGAGACTTCTTTCCCAGG |
| 2. osaA_R2 | CCCGAATTGATGGTGGACGTTGT |
| 3. osaA_F3 | GCACTCTTCCTACGATTACCCCGT |
| 4. osaa_R4 | ACCAAGCAGGATAAAGCTGATCGC |
| 5. osaA_F5 | ACAACGTCCACCATCAATTCGGGGGATCCTATGGATCTCAG |
| 6. osaA_R6 | ACGGGGTAATCGTAGGAAGAGTGCGTCGACATCACCCTTACCC |
| 7. osaA_F7 | CACTCCAGCACCAGTTGAACTGTTC |
| 8. osaA_R8 | CGGCGTTTTAACTGATGCACCTC |
| 9. A para pyrG_R | CAGGAGCAGCATAAATTCCACGACC |
| 10. osaA_com1 | AAGCTTAGTCATCTTCTCGCATGGCGC |
| 11. osaA_com2 | GCGGCCGCCATCAAGCGGCTCACATGACCA |
| 12. osaA_OE1 | AAAAAAAAGGCGCGCCATGGTGAACGGCACCGCT |
| 13. osaA_OE2 | AAAAAAAAGCGGCCGCTTACATTTGCGGCTGAGCTGGAG |
| 14. gpdA _F | AAGTACTTTGCTACATCCATACTCC |
| 15. trpC_R | TGCTTGATCTCGTCTCCCGAAA |
| 16. A fum pyrG_R | GAGCAGCGTAGATGCCTCGAC |
| 17. AfuTub-B_F | TTCCCAACAACATCCAGACC |
| 18. AfuTub-B_R | CGACGGAACATAGCAGTGAA |
| 19. His1 | CCGCCGTGGTGGTGTCAAG |
| 20. His2 | GGCGTGTTCAGTGTAGGTGACG |
| 21. osaA_CDS_F | CCGTGAGCTTGAAAAGCCGTTCC |
| 22. osaA_CDS_R | TCCACGTCCGACTGATGCGAT |
| 23. fumR_F | CCCAAGACCTCCTCGATCCAGTCCC |
| 24. fumR_R | GGTAGAAAGCCCATCACCAGCATCG |
| 25. pdsA_F | CGCAATACCCTGCACCGACTG |
| 26. pdsA_R | GTTTCGATCTGGAACAGAGCTTCGTTGA |
| 27. Afum_fmqD_F | ACGCTTGGGGGAGGGTTG |
| 28. Afum_fmqD_R | AAGTTCGCACCAGCACCCC |
| 29. gliZ_F_qRT | ACGACGATGAGGAATCGAACCCG |
| 30. gliZ_R_qRT | GGTGCTCCAGAAAAGGGAGTCGTTG |
| 31. gliP_F_qRT | AGTTACACCGACTCGCATCCAGC |
| 32. gliP_R_qRT | CTGGGGCAGACCATGCGTAG |
| 33. Afum_pypC_F | CATCGCACTCGCCTACTCCTTTCA |
| 34. Afum_pypC_R | GCAGTCTTGCTTTGACCGCAT |
| 35. Afum_psoA_F | CTCTGGCGGCGAGATTGGTT |
| 36. Afum_psoA_R | CCGCCCTTCTTTCCATCCTTCC |
| 37. AfumveA_F_new | TGCACCAATATCCGCCAATGC |
| 38. AfumveA_R_new | CGTGTAGGAAGGAGGAGGGGTTACC |
| 39. Afu_brlA_F | GTCAGCAAAGCCGAAGATGGACTACC |
| 40. Afu_brlA_R | GTTGTCGTGGCTCAAGGCGTAC |
| 41. Afum_qPCR_abaA_F | TTGACTCGCTGTCAAGTGATTGTGG |
| 42. Afum_qPCR_abaA_R | TGGTGGATTTGTGGTGGGGAGTT |
| 43. Afu_wetA_F | CTCTACTCACAGCTCTCATCGGGC |
| 44. Afu_wetA_R | GCTGCCATCGTGACTTACTGCG |
| 45. Afu_laeA_F | GTTGCCGTGCGACGAGCAGGA |
| 46. Afu_laeA_R | GGGGCCAAATCAACCCCAACGACG |
